# MiT4SL: multi-omics triplet representation learning for cancer cell line-adapted prediction of synthetic lethality

**DOI:** 10.1101/2025.04.20.649694

**Authors:** Siyu Tao, Yimiao Feng, Yang Yang, Min Wu, Jie Zheng

## Abstract

Synthetic lethality (SL) offers a promising approach for targeted cancer therapies. Current SL prediction models heavily rely on extensive labeled data for specific cell lines to accurately identify SL pairs. However, a major limitation is the scarcity of SL labels across most cell lines, which makes it challenging to predict SL pairs for target cell lines with limited or even no available labels in real-world scenarios. Furthermore, gene interactions could be opposite between training and test cell lines, i.e. SL vs. non-SL, which further aggravates the challenge of generalization among cell lines. A promising strategy is to transfer knowledge learned from cell lines with relatively abundant SL labels to those with limited SL labels for the discovery of novel SL pairs, i.e., cell line-adapted SL prediction. Here, we propose MiT4SL, a multi-omics triplet representation learning model for cell line-adapted SL prediction. The core idea of MiT4SL is to model cell lineage information as embeddings, which are generated by combining a protein-protein interaction network representation tailored to each cell line with the corresponding protein sequence embeddings. We then combine these cell line embeddings with gene pair representations derived from a biomedical knowledge graph and protein sequences. This triplet representation learning strategy enables MiT4SL to capture both shared biological mechanisms across cell lines and those unique to each cell line, effectively mitigating distribution shift and improving generalization to target cell lines. Additionally, explicit cell line embeddings provide the necessary signals for MiT4SL to effectively differentiate between cell line contexts, enabling it to adjust predictions and mitigate possible label conflicts for the same gene pair across different cell lines. Experimental results across various cell line-adapted scenarios show that MiT4SL outperforms six state-of-the-art models. To the best of our knowledge, MiT4SL is the first deep learning model designed specifically for cancer cell line-adapted SL prediction.

**Availability:** The code of our work is available at https://github.com/JieZheng-ShanghaiTech/MiT4SL.

## 1 Introduction

Synthetic lethality (SL) occurs between two genes when the perturbation of either gene alone is well tolerated, but their combined perturbation results in cell death [1]. Since gene mutation is a common feature of tumor cells, targeting SL partners of genes with cancer-specific mutations can selectively kill tumor cells while keeping normal cells alive (Figure 1A). A well-studied example of targeted therapy leveraging the concept of SL is the poly-ADP ribose polymerase (PARP) inhibitors targeting *BRCA-1/2* deficiency in the breast and ovarian cancers [2]. This successful case has sparked a gold rush in identifying new SL gene pairs as anti-cancer drug targets.

**Fig. 1.**
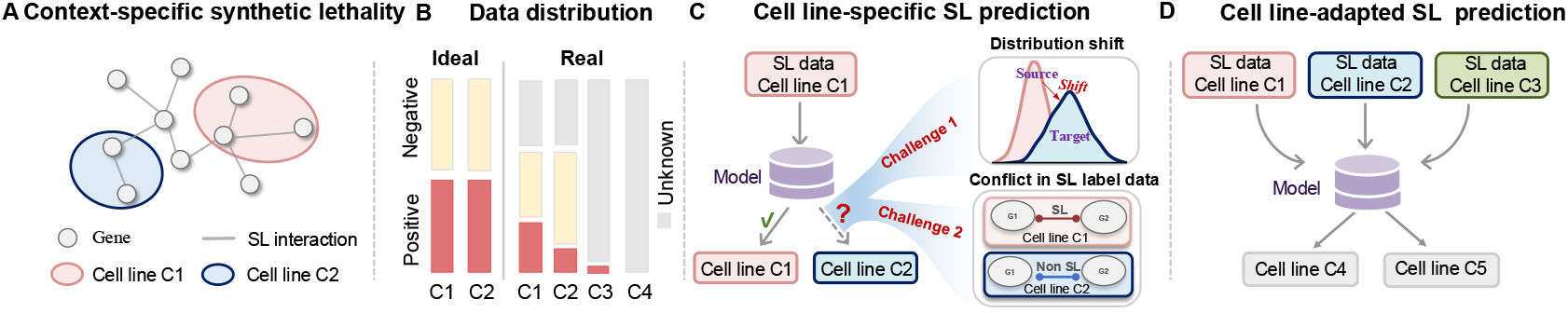
(A) The concept of context-specific synthetic lethality. (B) An ideal distribution (left). Real-world dataset distributions are often imbalanced, and many SL interactions are yet to be discovered in many cell lines (right). (C) Supervised learning models cannot be trained well due to the lack of SL label data in target cell lines. Two main generalization challenges exist for existing SL prediction methods when transferring to a target cell line: distribution shift and conflict of SL labels for the same pair of genes across different cell lines. (D) Illustration of the cell line-adapted SL prediction.

Experimental screening techniques, such as RNA interference [3] and CRISPR-based gene perturbation [4], have been developed to screen novel SLs. The experimental screening techniques are effective but have notably high costs and limitations, such as off-target effects, hindering their applications to large-scale gene search. By contrast, computational predictions can quickly narrow down the search scope of SL candidates, and thereby reduce the cost and time for SL screening. Therefore, machine learning methods for SL prediction have gained increasing attention and made much progress in the last few years [5]. Recently, a plethora of methods based on graph neural networks (GNNs) have been developed to predict potential SL gene pairs. These methods can be broadly divided into two main categories: context-free methods [6–10], which generate unified gene representations to predict SLs without considering genetic context, and context-specific approaches [11–14], which identify SLs by learning gene representations from the genetic background of individual cancer types or cell lines.

Despite such progress, a main challenge in the field is the scarcity of available labeled data (Figure 1B). To date, out of 1000 established cancer cell lines in the Cancer Cell Line Encyclopedia (CCLE) database [15], only 31 cell lines (3.1%) have SL labels available. Additionally, in each of these 31 cell lines, only a tiny fraction of the hundreds of millions of gene pairs have known SL interactions. Therefore, in real-world applications, SL candidate pairs need to be predicted for those cell lines where SL labels are severely scarce or even completely absent. Faced with the data sparsity issue, existing SL prediction methods struggle to be effectively trained. Due to their inherent context-specificity, SL gene pairs in cell lines to be explored (i.e. the target cell lines) are often unseen or differ significantly from those in source cell lines used as training data, leading to the issue of distribution shift (Figure 1C). Unfortunately, the limited SL labels available in a single cell line impose a narrow scope of information for predictive models, which could severely hinder their generalizability to target cell lines. Furthermore, recent studies [16–18] have shown that SL interactions between the same pair of genes could exhibit variations or conflicts across different cell lines (Figure 1C).

For instance, gene *ARIH1* is synthetic lethal with gene *TIPIN* in the Jurkat cell line, but not so in the K562 cell line [17]. Such conflicts of SL labels across cell lines introduce another challenge for generalization.

Recently, in the field of transfer learning, *domain generalization*, a strategy of learning generalizable representations from diverse source domains to address label data sparsity in the target domain, has achieved success in tasks such as image classification and natural language processing [19–21]. Inspired by these advancements, we propose that a promising strategy for addressing the scarcity and conflict of SL label data across cell lines is to develop robust models capable of transferring knowledge from cell lines with relatively abundant SL labels to target cell lines, which we formulate as the task of cell line-adapted SL prediction (Figure 1D).

In this paper, we propose MiT4SL, a novel deep learning model that integrates three-way (i.e. triplet) representations of genes and cell lines learned from multi-omics data, to tackle the challenge of data sparsity in cell line-adapted SL prediction. Specifically, MiT4SL encodes cell line information as independent embeddings, enabling awareness of cell line context. By identifying cell line-specific subgraphs of a protein-protein interaction (PPI) network using gene expression data, and incorporating context-free gene representations learned from a biological knowledge graph and the protein language model of ESM-2, MiT4SL is able to capture signals about biological mechanisms shared by various cell lines as well as those unique to each cell line. As such, MiT4SL can alleviate the biases and noise caused by the aforementioned distribution shift and label conflict across cell lines. Extensive experiments demonstrate the superior performance of MiT4SL against six state-of-the-art (SOTA) baselines. Case studies also confirm the potential of MiT4SL in predicting promising novel SLs. In conclusion, to address the challenges due to scarcity and noise of SL label data, we propose a cell line-adapted strategy for SL prediction, and a novel method MiT4SL with promising accuracy and generalizability. Our work would pave the way for mining in an unexplored landscape of novel anti-cancer drug targets.

## 2 Methodology

### 2.1 Problem formulation

Let 𝒢 = *{g*_1_, *g*_2_, *· · ·*, *g*_*N*_ *}* and *C* = *{*c_1_, c_2_, *· · ·*, c_*K*_*}* represent sets of genes and cell lines. We define the training cell lines as 𝒞_train_ ⊂ 𝒞, and the test cell lines as 𝒞_test_ ⊂ 𝒞 *\* 𝒞_train_. Furthermore, the gene sets of the cell lines 𝒞_train_ and 𝒞_test_ are denoted by 𝒢 _train_, 𝒢 _test_ ⊂ 𝒢, respectively. In this study, we focus on a cell line-adapted SL prediction problem: *Given the gene pairs* (*g*_*i*_, *g*_*j*_) ∈ *𝒢*_*train*_ *in the known cell lines* 𝒞_*train*_, *we aim to construct a model to predict the SL relationships of gene pairs* 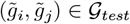 *in the unknown cell lines 𝒞*_*test*_.

To address this problem, we propose to construct a set of triplets 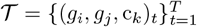 from *G*_train_ and 𝒞_train_, where *g*_*i*_ and *g*_*j*_ are two different genes in 𝒢_train_ and c_*k*_ is a cell line in 𝒞_train_. We then introduce a multi-omics triplet representation learning model to learn a scoring function ℱ: (*g*_*i*_, *g*_*j*_, c) →*ŷ* that estimates the probability of a SL relationship between the gene pair (*g*_*i*_, *g*_*j*_) in the training cell line c_*k*_. Here, *ŷ* ∈ [0, 1] is the predicted score, with a higher value indicating a stronger likelihood of an SL relationship between *g*_*i*_ and *g*_*j*_ in c_*k*_. After training, we apply the model to make predictions for gene pairs 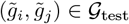 in target cell lines c_ℓ_∈𝒞 _test_. Specifically, for each 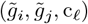 in the test set, the model outputs a predicted score *ŷ*, indicating the probability of a SL relationship between 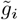 and 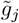 in c_ℓ_.

### 2.2 Overview of MiT4SL

Figure 2 shows the overview of the proposed MiT4SL model. Given the *t*-th input triplet (*g*_*i*_, *g*_*j*_, c_*k*_)_*t*_, we firstly use a Biomedical knowledge graph (BKG) encoder ℬ_*ϕ*_ (Sec. 2.3) and a protein sequence (PS) encoder 𝒫_*φ*_ (Sec. 2.4) to extract the BKG-based representations {**z**_kg_(*g*_*i*_), **z**_kg_(*g*_*j*_)} and sequence-based representations {**z**_seq_(*g*_*i*_), **z**_seq_(*g*_*j*_)} for the gene pair (*g*_*i*_, *g*_*j*_). Meanwhile, we use transcripts per million (TPM) to determine highly expressed genes for the input cell line c_*k*_ from to construct a contextual 𝒢 sub-PPI network specific to the cell line c_*k*_ from a PPI network. Then, we adopt a cell line encoding strategy (Sec. 2.5) to produce the cell line representation **h**(c_*k*_) from the sub-PPI network and the protein sequence-based representations. Finally, the triplet representation (*g*_*i*_, *g*_*j*_, c_*k*_)_*t*_ can be generated by incorporating the above multi-omics representations. To optimize our model (Sec. 2.6), we feed the triplet representation into the SL predictor to estimate the probability of SL relationship between the gene pair (*g*_*i*_, *g*_*j*_) in the cell line c_*k*_ and compute two novel regularizations to enhance the triplet representation.

**Fig. 2.**
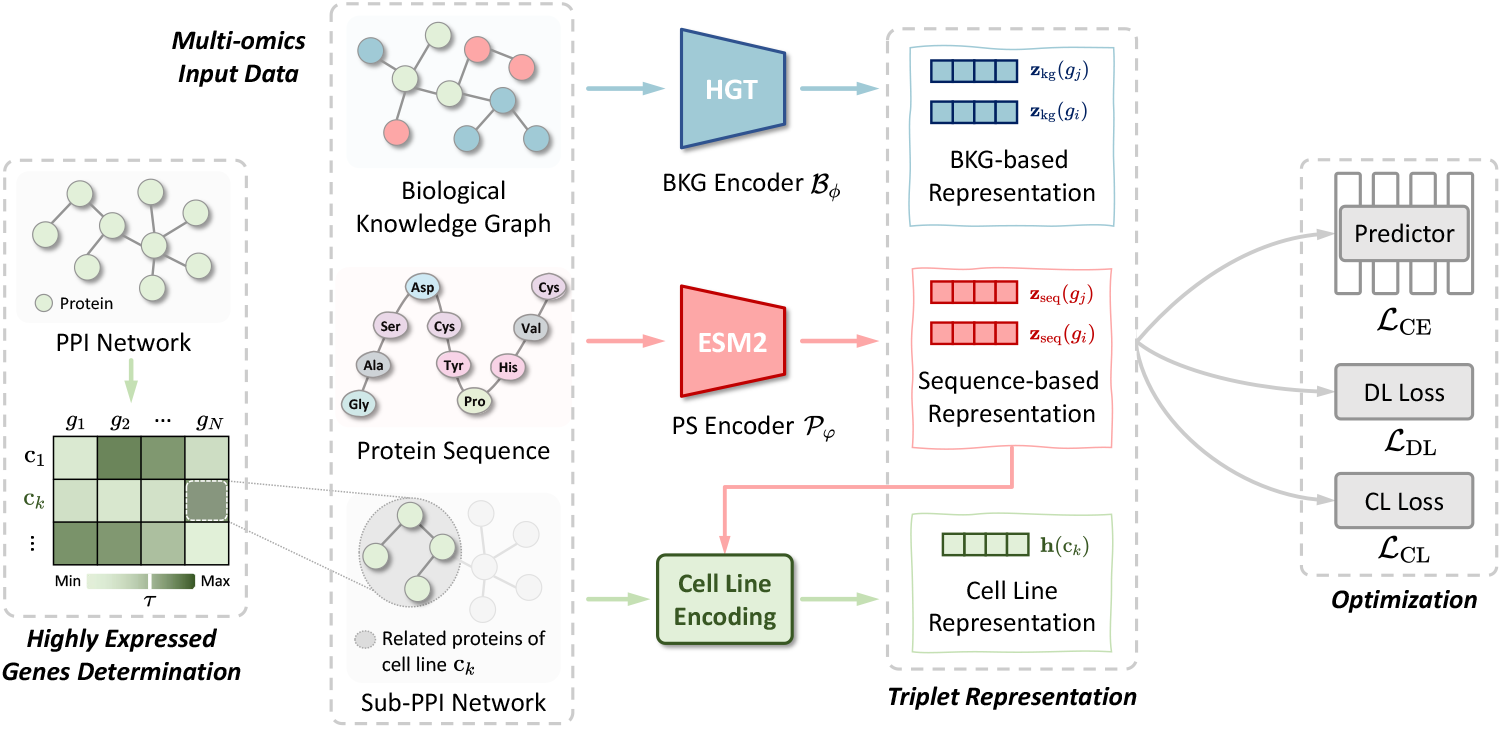
Overview of the proposed MiT4SL model. Firstly, we feed a biomedical knowledge graph and protein sequences into a BKG encoder and a PS encoder to generate BKG-based and sequence-based representations for input gene pairs. Meanwhile, we use TPM to determine highly expressed genes for the cell line and produce a cell line-specific sub-PPI network. Then, we conduct our cell line encoding to produce the cell line representation from the sequence-based representations and the sub-PPI network. The triplet representation is thus yielded by concatenating these multi-omics representations. Finally, we optimize our MiT4SL by minimizing predicted loss on SL scores and two regularizations to enhance the triplet representation.

### 2.3 Biomedical knowledge graph encoder

Through integrating diverse data sources and capturing complex biological interactions among biological entities [8, 22, 23], BKGs have the potential to generate effective gene representations, thereby enhancing SL prediction performance. Consequently, to extract the semantic interactions from a BKG, we introduce a BKG encoder to perform BKG-based representation learning for each gene *g*. Specifically, a BKG is represented as 𝒦 𝒢= (𝒱, ℰ), where 𝒱 and ℰ are sets of nodes and edges, respectively. In addition, each pair of nodes in the BKG is denoted by a triplet (*v*_*i*_, *v*_*j*_, *e*_*ij*_), where *v*_*i*_, *v* _*j*_∈𝒱 and *e*_*ij*_∈ℰ. A triplet in the BKG 𝒦 𝒢 contains a certain kind of semantic interaction between the two biological entities. For instance, the triplet (*TP53, DNA damage, interacts with*) means that the gene *TP53* interacts with the pathways of DNA damage responses. Generally, the learning process can be expressed as:

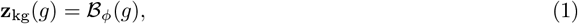

where ℬ_*ϕ*_ is the BKG encoder, *ϕ* represents its trainable parameters, and **z**_kg_(*g*) ∈ ℝ^*D*^ is the BKG-based representation of gene *g*.

In MiT4SL, we adopt the Heterogeneous graph transformer (HGT), a GNN model designed for the representation learning of general heterogeneous graphs [24], to implement the BKG encoder ℬ_*ϕ*_. Specifically, we first use the efficient graph sampling strategy provided by HGT to divide the large-scale BKG *KG* into a set of small sub-graphs to reduce memory usage. Then, the BKG-based representation 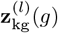 in the *l*-th HGT convolutional layer is generated by three operators: heterogeneous mutual attention, heterogeneous message passing, and target-specific aggregation. Formally, this process can be summarized as:

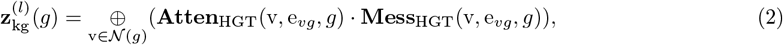

where 𝒩 (*g*) is the set of neighbors of gene *g* in the BKG, e_*vg*_ is the edge between gene *g* and its neighbor node v ∈ 𝒩 (*g*), **Atten**_HGT(*·*)_ is the heterogeneous mutual attention estimating the importance of the neighbor v, **Mess**_HGT_(*·*) is the heterogeneous message passing that extracts the messages from the neighbor v, and ⊕ (*·*) is the target-specific aggregation that averages the messages from all the neighbors. Besides, we use Xavier’s uniform distribution to initialize the representation 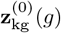,where nodes belonging to the same category have the same initial representation. Finally, we apply a non-linear activation function **GELU** followed by a type-specific linear projection **Line**_*τ*(*g*)_ and a residual connection 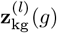,where *τ* (*g*) denotes the node type of gene *g* in the BKG. Finally, the BKG-based representation of gene *g* can be written as:

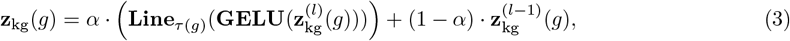

where *α* is a learnable parameter that adaptively determines the relative importance of the two components. In this study, we use the PrimeKG [25] as the BKG.

### 2.4 Protein sequence encoder

While the BKG provides interaction information between biological entities, protein sequences contain rich biological functional and structural information, which is crucial to identify SLs in terms of biological function perspective [9, 13, 26]. As such, we believe that incorporating interaction information from the BKGs and protein sequences could be an effective way to produce comprehensive gene representations and thus enhance the identification of SL pairs. To this end, we introduce a protein sequence (PS) encoder to generate the protein-sequence-based representation for each gene *g*. Generally, this process can be summarized as:

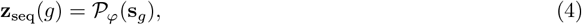

where 𝒫_*φ*_ is the PS encoder, *φ* denotes its learnable weights, **s**_*g*_ is the protein sequence, and **z**_seq_(*g*) ∈ ℝ^*D*^ represents the sequence-based representation.

In MiT4SL, we employ ESM2 [26] followed by a 2-layer MLP network to implement the encoder 𝒫_*φ*_. The ESM2 is a cutting-edge pre-trained protein language model for encoding protein sequences. It is pre-trained on the UniRef database and the protein sequences **s**_*g*_ are taken from the UniProt database [27]. In this study, we freeze the ESM2 model and only optimize the MLP network. Protein sequences are obtained from UniProt database.

### 2.5 Encoding cell lines with multi-omics data

Generating effective and generalizable representations of cell lines plays an indispensable role in cell line-adapted SL prediction due to the genetic background diversity of cell lines. As such, to create comprehensive cell line representations, our MiT4SL model adopts a novel cell line encoding strategy that incorporates graph representations from each cell line’s contextual PPI network with corresponding protein sequence embeddings from the ESM2 model 2.4. To this end, given a cell line c ∈ *𝒞*, we first identify a cell line-specific gene set 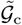 from *𝒢*. This process includes three steps: 1) based on TPM provided by [28], we set a threshold *τ* to determine a highly expressed gene set 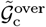 from *G*; 2) for each highly expressed gene 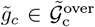,we sample its *q* neighbor nodes in a PPI network [29] to form 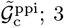) finally, the gene set 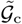 specific to the cell line c is constructed by 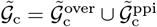.The representation learning for the cell line c ∈ *𝒞* can thus be transformed into encoding the cell line-specific gene set 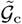.

PPI network data have been shown to effectively reflect contextual information [30, 31]. To this end, we first extract a sub-PPI network tailored to the input cell line from the full PPI network, based on the cell line-specific gene set 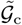. Then, two modalities of data (sub-PPI network and protein sequences) are used to construct the encoding, which can effectively enrich the cell line representation. Specifically, we first use Xavier’s uniform distribution to initialize the representation of each node in the sub-PPI network. Then, we feed the graph into a GNN model DeeperGCN [32] to obtain the PPI-based representation 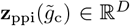, where 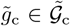.Meanwhile, we feed the protein sequences of these genes to the PS encoder *P*_*φ*_ to generate the corresponding sequence-based representation 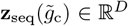.The two components of a gene 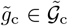 are defined:

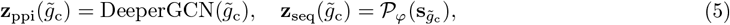

where 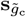is the protein sequence of the gene 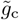.Then, using a pooling mechanism, the two modalities representations **h**_ppi_(c) ∈ ℝ^*D*^ and **h**_seq_(c) ∈ ℝ^*D*^ of the cell line c are generated by:

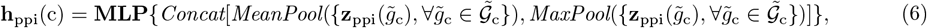

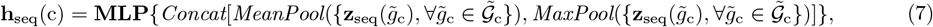

where **MLP** are two single-layer MLP networks. Moreover, *MaxPool, MeanPool*, and *Concat* denote the max-pooling operator, mean-pooling operator, and concatenation operator, respectively. Finally, the cell line representation **h**(c) ∈ ℝ^2*D*^ is defined as:

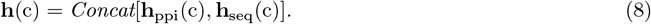

As such, the cell line encoding is completed.

### 2.6 Optimization

To optimize learnable components, including the BKG encoder, PS encoder, DeeperGCN, and MLP networks, in the MiT4SL model, we use the back-propagation algorithm to minimize the following three losses.

#### Synthetic lethality prediction

We first compute a binary cross-entropy loss ℒ_CE_ that measures the distance between predicted and real SL scores. Specifically, for the *t*-th input triplet (*g*_*i*_, *g*_*j*_, c_*k*_)_*t*_, we first use the above encoders to generate its multi-omics representation {**z**_kg_(*g*_*i*_), **z**_kg_(*g*_*j*_), **z**_seq_(*g*_*i*_), **z**_seq_(*g*_*j*_), **h**(c_*k*_)}. Then, a 3-layer MLP classifier takes the triplet representation as input and then predicts the SL score *ŷ*_*t*_. This process can be expressed as:

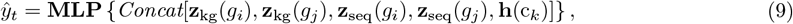

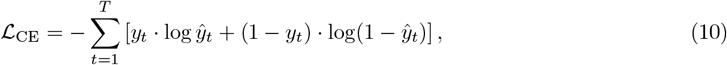

where **MLP** represents the classifier with hidden sizes of [128, 128, 64] and *y*_*t*_ is the real SL score.

#### Enhancing triplet representations via contrastive learning

In MiT4SL, the entire multi-omics representation of a triplet (*g*_*i*_, *g*_*j*_, c_*k*_)_*t*_ includes five items **z**_kg_(*g*_*i*_), **z**_kg_(*g*_*j*_), **z**_seq_(*g*_*i*_), **z**_seq_(*g*_*j*_), and **h**(c_*k*_) from four types of biological data (i.e., a BKG, protein sequences, gene expression data, and a PPI network). We propose a contrastive learning-based regularization to integrate these data, thereby improving effectiveness of the representations. Specifically, there are four combinations of embeddings of two genes, as each gene has two types of embeddings. We thus divide the triplet representation into four distinct sub-representations, defined by: **Z**_*t*_(*u, v*) = {**z**_*u*_(*g*_*i*_), **z**_*v*_(*g*_*j*_), **h**(c_*k*_)}, *u, v* ∈ *{*seq, kg}. For example, a sub-representation **Z**_*t*_(kg, seq) denotes the combination of the BKG-based embedding of the gene *g*_*i*_, the sequence-based embedding of the gene *g*_*j*_, and the cell line embedding **h**(c_*k*_). Then, by considering positive pairs as two sub-representations from the same triplet, and negative pairs as those from different triplets, we define a contrastive learning loss ℒ_CL_:

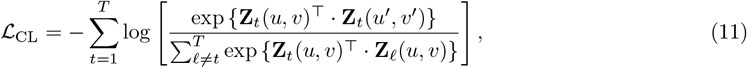

where **Z**_*t*_(*u, v*) and **Z**_*t*_(*u*^*′*^, *v*^*′*^) are any two different sub-representations from the same triplet (*g*_*i*_, *g*_*j*_, c_*k*_)_*t*_, such as **Z**_*t*_(kg, kg) and **Z**_*t*_(kg, seq). The goal of the regularization ℒ_CL_ is to minimize the difference of sub-representations from the same triplet, while maximizing the difference of those from different triplets.

Therefore, we can effectively integrate the features of the multi-omics data, which can significantly improve the SL prediction performance.

#### Decision-level regularization

Improving the consistency of the sub-representations from the same triplet can enhance the generalizability of the entire triplet representation. Here, a decision-level regularization ℒ_DL_ is proposed to further enhance this consistency. Specifically, we feed the BKG-based sub-representation **Z**_*t*_(kg, kg) and sequence-based sub-representation **Z**_*t*_(seq, seq) from the same triplet (*g*_*i*_, *g*_*j*_, c_*k*_)_*t*_ into an MLP classifier to predict SLs. The process is summarized as follows:

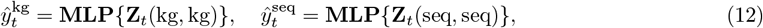

where **MLP** denotes the MLP classifier. 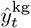 and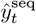 are the predicted SL scores corresponding to the sub-representations **Z**_*t*_(kg, kg) and **Z**_*t*_(seq, seq). To maximize the consistency between **Z**_*t*_(kg, kg) and **Z**_*t*_(seq, seq), we average the two scores to generate the final SL score. The regularization *L*_DL_ is thus defined as:

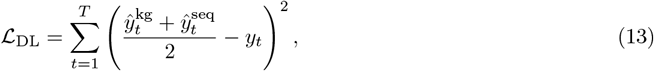

where *y*_*t*_ is the real SL label for the triplet (*g*_*i*_, *g*_*j*_, c_*k*_)_*t*_.

#### Overall loss

In summary, our full loss function ℒ_Overall_ is defined as:

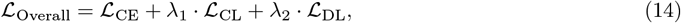

where *λ*_1_ *>* 0 and *λ*_2_ *>* 0 are two hyper-parameters that control the contributions of the regularizations ℒ_CL_ and ℒ_DL_. Their effects on the model performance are described in the Supplementary Material.

## 3 Experimental settings

### 3.1 Datasets

#### Synthetic lethality datasets

We construct the synthetic lethality (SL) dataset based on the genetic interaction scores (GI scores) provided by a public SL dataset named SLKB [33]. For each pair of genes, the GI score evaluates their functional relationship by quantifying the difference between the observed phenotypic effect of their co-mutations and the expected effect derived from their individual mutations. We formulated the samples by following the rule in [17]: each quadruple (Gene A, Gene B, cell line C, GI score) in the original dataset is viewed as a positive SL pair if the GI score is below − 3; otherwise, it is viewed as a negative (i.e., non-SL) pair. Finally, we obtained 6,883 gene pairs from 6 widely-used types of cell lines, i.e., A375, A549, Jurkat, Mewo, 22Rv1, and Pk1. Statistics of the SL dataset are provided in Table 1. This is a highly imbalanced dataset, where only 1,361 samples are SL pairs. We constructed the cell line-adapted evaluation scenarios by performing the leave-one-cell-line-out data splitting. Specifically, in the cell line-adapted setting, the gene pairs from multiple cell lines are used for model training, and then the trained models are tested on a target cell line not included in the training data. Furthermore, to avoid data leakage, we removed from the input features those edges in the BKG [25] and PPI network [29] overlapping with the gene pairs labeled as SL or non-SL in SLKB.

**Table 1.**
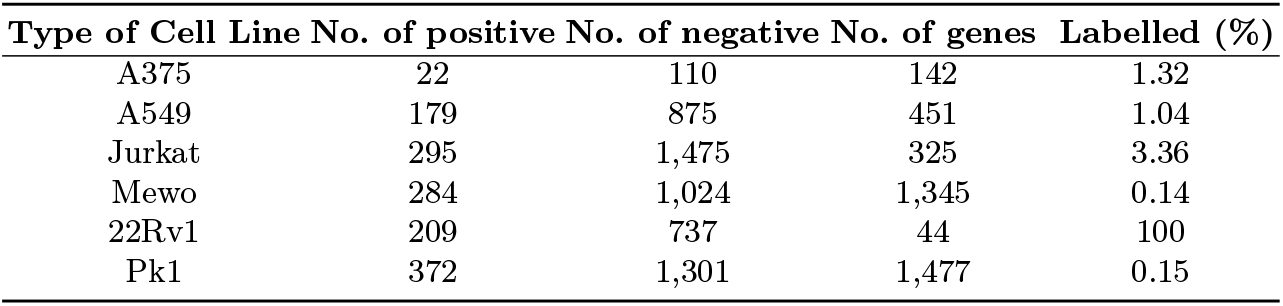
Statistics of six cell lines in our SL dataset. “Labelled (%)” denotes the percentage of labelled SL gene pairs out of all possible gene pairs involving genes from a cell line.

#### Multi-omics feature datasets

We construct effective triplet representations using four types of biological features: a biomedical knowledge graph PrimeKG [25], protein sequences from the UniProt database [27], a protein-protein interaction (PPI) network from a previous study [29], and transcriptomic data from the Sanger database [28]. A summary of the multi-omics datasets used in our study is provided in Table 2.

**Table 2.** Summary of the four multi-omics feature datasets.

| Feature | Data information | Source |
| --- | --- | --- |
| Biomedical knowledge graph | 129,075 nodes and 8,100,498 edges | PrimeKG [25] |
| Protein sequences | 19,015 protein sequences | UniProt [27] |
| PPI network | 17,161 proteins and 839,522 edges | Titeca.*et al* [29] |
| Transcriptomic data | 6 cell lines × 37,263 genes | Sanger [28] |

### 3.2 Baselines

We compare MiT4SL against six SOTA methods for SL prediction, comprising four cell line-free methods [7–10] and two cell line-specific methods [12, 14]. The cell line-free models, i.e. KG4SL [8], PTGNN [9], NSF4SL [10], and SLMGAE [7], predict SLs without using cell line information. By contrast, SLGNNCT [14] and MVGCN-iSL [12] leverage cell line information to identify SLs. An introduction to the baseline models is given in the Supplementary Material. We ran the baseline models using the code provided by the authors to ensure a fair comparison.

### 3.3 Implementation details

In MiT4SL, we use HGT [24] to implement the BKG encoder ℬ. It consists of 4 layers of the HGT blocks, where the hidden sizes of the layers are set to [256, 128], and the number of attention heads is set to 4. Since PrimeKG is a large knowledge graph, we use a sampling strategy from [24] with four sampling layers to collect subgraphs related to input genes. For each input gene, each sampling layer samples 1024 nodes. The dimension of the gene features is 64 (i.e., *D* = 64). We use the Adam optimizer with default hyperparameters. In the cell line-adapted scenario, due to significant variation in training data across different target cell lines, a range of different learning rates and epoch parameters is used. Specifically, the learning rate varies between 0.001 and 0.002, with epoch counts of 20, 50, and 80. The weights of the two regulations in the overall loss, *λ*_1_ and *λ*_2_, are both set to 0.2. Moreover, we set the number of sampled neighbor nodes *q* = 32 in the PPI network. Following [30], the threshold of TPM *τ* is set to 400.

## 4 Results and Discussions

### 4.1 Performance comparison in the cell line-adapted SL prediction

To systematically evaluate the effectiveness of MiT4SL in SL predictions, we construct six cell line-adapted evaluation combinations. For example, the gene pairs from the A375 cell line serve as test data, while the remaining five cell lines are used for training. Note that MVGCN-iSL [12] cannot be applied to cell line-adapted SL predictions, as it allows for only a single cell line in the training or test data.

Our initial observation is that MiT4SL achieves the best overall performance compared to all baselines (Table 3 and Table 4). For example, MiT4SL outperforms the second-best baseline SLGNNCT by 8.71% in the AUC score and 5.05% in the BACC score, respectively. More importantly, we find that MiT4SL achieves substantial improvements in specific cell lines where baseline models struggle to perform effectively. For instance, although SLMGAE has been benchmarked as a SOTA SL prediction method [5], it achieves only an AUC score of 0.4270 in the A549 cell line. In comparison, MiT4SL achieves an AUC score of 0.6366, marking a 20.96% improvement over SLMGAE. Similarly, when 22Rv1 is the target cell line, MiT4SL significantly outperforms all baselines, achieving an AUC score increase of at least 15.05%. These promising results underscore MiT4SL’s strong adaptability to diverse biological contexts, allowing it to generalize effectively to target cell lines. Secondly, MiT4SL consistently outperforms all baselines in terms of BACC scores across various cell line-adapted evaluations. By comparison, some baselines, including but not limited to KG4SL, often achieve a BACC score of 0.5, indicating a tendency to classify all gene pairs as negative. These results may be attributed to the fact that these models are biased toward the majority negative class in the training data, highlighting their weak robustness in handling imbalanced SL predictions.

**Table 3.** Performance comparison between MiT4SL and existing approaches across six evaluation combinations for the cell-line-adapted SL prediction task. The best and runner-up performances (in AUC scores) are highlighted in **bold** and underline, respectively.

| Model | Target Cell Line |  |  |  |  |  | Avg. |
| --- | --- | --- | --- | --- | --- | --- | --- |
|  | A375 | A549 | Jurkat | Mewo | 22Rv1 | Pk1 |  |
| SLMGAE [7] | 0.6568±0.02 | 0.4270±0.00 | <u>0.6148±0.00</u> | 0.6075±0.00 | 0.3958±0.01 | <u>0.6418±0.00</u> | 0.5573 |
| KG4SL [8] | 0.4359±0.02 | 0.4898±0.03 | 0.5181±0.05 | 0.5413±0.02 | <u>0.4632±0.05</u> | 0.5611±0.01 | 0.5016 |
| PTGNN [9] | 0.4894±0.02 | 0.4332±0.02 | 0.4845±0.02 | 0.6410±0.04 | 0.3985±0.01 | 0.6035±0.01 | 0.5084 |
| NSF4SL [10] | 0.6608±0.10 | 0.5008±0.05 | 0.5434±0.03 | 0.5865±0.02 | 0.4490±0.03 | 0.5324±0.02 | 0.5455 |
| SLGNNCT [14] | 0.5551±0.04 | <u>0.5775±0.04</u> | 0.4830±0.05 | <u>0.6441±0.04</u> | 0.4557±0.06 | 0.6393±0.03 | <u>0.5591</u> |
| MiT4SL (Ours) | **0.6781±0.05** | **0.6366±0.03** | **0.6200±0.00** | **0.6757±0.00** | **0.6137±0.02** | **0.6532±0.00** | **0.6462** |

**Table 4.** Performance comparison of MiT4SL and existing approaches across six evaluation combinations for the cell-line-adapted SL prediction task. The best and runner-up performances (in BACC scores) are highlighted in **bold** and underline, respectively.

| Model | Target Cell Line |  |  |  |  |  | Avg. |
| --- | --- | --- | --- | --- | --- | --- | --- |
|  | A375 | A549 | Jurkat | Mewo | 22Rv1 | Pk1 |  |
| SLMGAE [7] | 0.5834±0.02 | 0.4234±0.00 | 0.5000±0.00 | 0.5127±0.03 | 0.4525±0.01 | <u>0.6135±0.00</u> | 0.5143 |
| KG4SL [8] | 0.5000±0.00 | 0.5000±0.00 | 0.5000±0.00 | 0.5000±0.00 | 0.5000±0.00 | 0.5000±0.00 | 0.5000 |
| PTGNN [9] | 0.5232±0.01 | 0.4524±0.01 | 0.4964±0.00 | 0.4992±0.00 | 0.4999±0.01 | 0.5490±0.01 | 0.5034 |
| NSF4SL [10] | 0.5000±0.00 | 0.5000±0.00 | 0.5000±0.00 | 0.5000±0.00 | 0.5000±0.00 | 0.5000±0.00 | 0.5000 |
| SLGNNCT [14] | <u>0.6032±0.09</u> | <u>0.5422±0.01</u> | 0.4813±0.03 | <u>0.5827±0.03</u> | 0.4766±0.04 | 0.5785±0.03 | <u>0.5441</u> |
| MiT4SL(Ours) | **0.667±0.03** | **0.5827±0.03** | **0.5156±0.00** | **0.6354±0.01** | **0.5425±0.02** | **0.6243±0.00** | **0.5946** |

Given the high cost and low throughput of wet-lab validation, we next evaluate the effectiveness of MiT4SL in accurately recommending SL partners for selected primary genes. Specifically, we choose six key genes as primary genes in the 22Rv1 cell line, including *AR, AKT3*, and *EZH2*. Each primary gene has 43 candidate SL partners. We define the top 20% of predictions (43 × 20% ≈ 8) as the recommended results and evaluate performance using Precision@8, which ranges from 0 to 1. SLGNNCT [14] and SLMGAE [7], two best performance baselines, are included in this comparison. Based on the results, we highlight two significant findings (Fig 3). Firstly, MiT4SL achieves the highest Precision@8 across all primary genes, demonstrating its superiority in prioritizing relevant SL candidate pairs. Secondly, taking a bird’s-eye view of the full SL partner networks for primary genes, MiT4SL captures a more diverse range of SL interactions, as evidenced by the larger red clusters across all networks. Therefore, these findings highlight MiT4SL’s potential in prioritizing candidate gene pairs for experimental validation, which may help address the urgent need for cost-effective solutions in SL discovery. Additionally, results for other primary genes (*GART, MAPK13*, and *WNT5A*) are provided in the Supplementary Material.

**Fig. 3.**
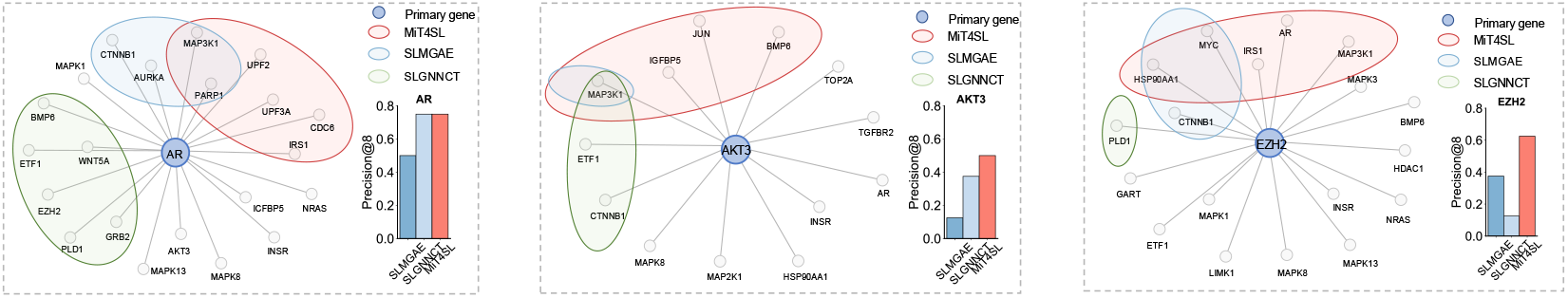
Performance comparison of MiT4SL and baseline models in recommending SL partners for primary genes in the 22Rv1 cell line. Quantitative results are evaluated using the Precision@8 score, which ranges from 0 to 1, with higher values indicating better performance. Nodes represent genes, with the primary gene highlighted in blue.

Furthermore, we evaluate MiT4SL in two additional SL prediction scenarios: cell line-specific and One2One. The cell line-specific scenario involves training and testing within the same cell line, whereas the One2One scenario trains on one cell line and transfers to another. For comparison, we focus on three baseline models, including SLMGAE, SLGNNCT [14], and MVGCN-iSL [12]. MiT4SL consistently outperforms other models across these scenarios (Table 5). Firstly, all models perform best in the cell line-specific scenario, as there is no biological background shift within the same cell line. Secondly, all models face challenges in the One2One scenario, likely due to the limitations of single-cell line data, which hinder their generalization ability. Nevertheless, MiT4SL demonstrates relatively better performance than the baselines. Full results are available in the Supplementary Material.

**Table 5.**
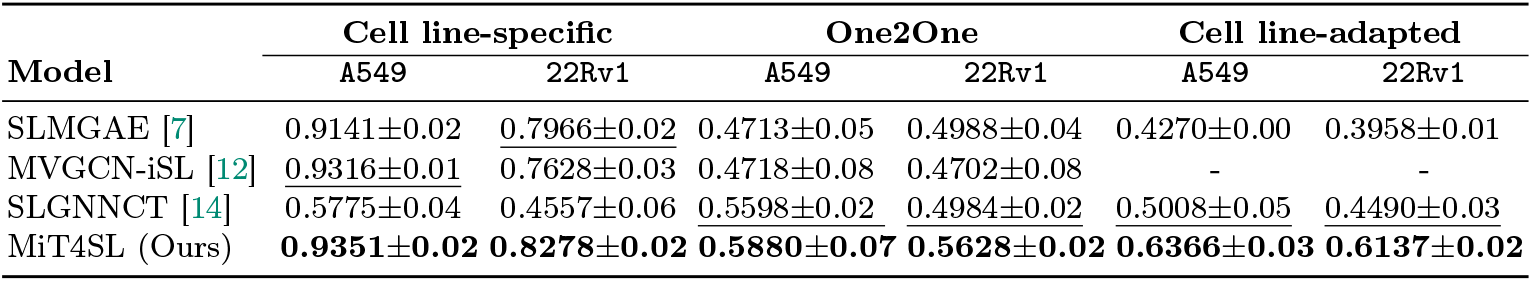
Performance comparison of MiT4SL and baselines across three evaluation scenarios (AUC score).

### 4.2 Impact of the cell line representation

Having validated the superior performance of MiT4SL, we next assess the role of cell line representations in mitigating distribution shift when generalizing to target cell lines, and their effectiveness in addressing SL label conflicts across different cell lines.

#### Mitigating distribution shift by introducing the cell line representations

To evaluate the importance of cell line representation in MiT4SL within the cell line-adapted scenario, we first conduct an ablation study by comparing the full MiT4SL model with MiT4SL (W/o Cell Line Repre.), which removes cell line representation and adopts a two-tuple representation. Other model configurations remain unchanged to ensure a fair comparison. Results show that incorporating cell line representation leads to a 4% increase in AUC score (Figure 4A), clearly demonstrating its significance in cell line-adapted scenarios.

**Fig. 4.**
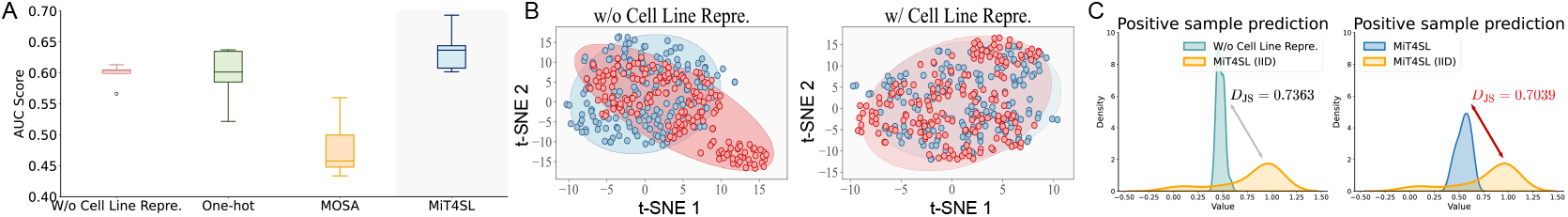
(A) Quantitative results of ablating the cell line representation in our MiT4SL for the cell line-adapted SL prediction (Target cell line A549). (B) Visualization of the embeddings for positive SL pairs with and without the cell line representation in MiT4SL for cell line-adapted SL prediction. (C) Visualization of prediction distributions with and without cell line representation in cell line-adapted scenarios, along with the prediction distribution in the cell line-specific scenario (denoted as IID). *D*_JS_ quantifies the distributional differences.

Then, we further explore whether introducing the cell line representations can truly enhance the generalizability of models. We visualize the embeddings of the positive SL pairs by ablating the cell line representation. Specifically, we map the high-dimensional embeddings of the positive SL pairs from training data and test data (A549) into a 2D space by using t-SNE [34]. Compared to MiT4SL (W/o Cell Line Repre.), the full MiT4SL (W/ Cell Line Repre.) can better align novel test positive SL pairs to the learned positive SL pairs manifold (Figure 4B). Moreover, we find that incorporating cell line representation improves the alignment of the predicted SL score distribution with the ideal distribution observed in the cell line-specific (IID) setting (Figure 4C). This is evidenced by a lower Jensen-Shannon (JS) divergence [35], denoted as *D*_*JS*_, where a smaller value indicates greater distributional consistency with the IID scenario. Full analysis of this result refers to Supplementary Material. These results illustrate that explicit cell line information as representations could aid the MiT4SL in capturing biological mechanisms shared by diverse cell lines, which enables it to better generalize to target cell lines.

Next, we compare MiT4SL against two alternative cell line encoding strategies to assess the effectiveness of its encoding approach. Specifically, MiT4SL (One-hot), which applies a simple one-hot encoding, and MiT4SL (MOSA), which leverages a SOTA cell line representation method [36]. MiT4SL achieves the best performance (Figure 4A), verifying the effectiveness of its cell line encoding strategy. Moreover, we analyze both feature types in the cell line representations in the Supplementary Material and find that each contributes meaningfully, supporting the validity of their integration.

In conclusion, the cell line representation learning strategy adopted in MiT4SL enables the model to effectively capture shared biological mechanisms across cell lines. These shared biological mechanisms could serve as the bridge to mitigate the distribution shift from source to target cell lines.

#### Alleviating the conflicting of SL labels across various cell lines by introducing the cell line representations

Compared to One2One scenarios, both SLMAGE and SLGNNCT experience a significant performance drop in the cell line-adapted scenarios. For instance, the AUC score of SLGNNCT decreases by 5.9% when evaluated on the target cell line A549. By comparison, MiT4SL gains benefits when adding more training cell lines. This observation raises a critical question: Why does adding more cell lines to the training data improve MiT4SL while harm other models? To investigate this, we evaluate MiT4SL alongside the two best-performing baselines, SLMGAE and SLGNNCT, using A549 as the test cell line. Specifically, we start with Mewo as the training cell line and progressively add others, increasing the number of training cell lines from one to five. MiT4SL consistently outperforms all baselines across all scenarios and shows continuous performance gains as more cell lines are introduced. However, baseline models exhibit minimal improvement or even performance degradation in some cases (Figure 5A). After examining the training and test data, we find that 83 gene pairs have conflicting labels across cell lines, accounting for approximately 10% of the test data. For instance, the gene pair *MCL1-BCL2L1* is labeled as a positive SL pair in A549 (test data) but as a negative sample in the training data. Therefore, we hypothesize that conflicts in SL interactions across cell lines may serve as an obstacle, preventing models from effectively leveraging more labeled data from multiple cell lines.

**Fig. 5.**
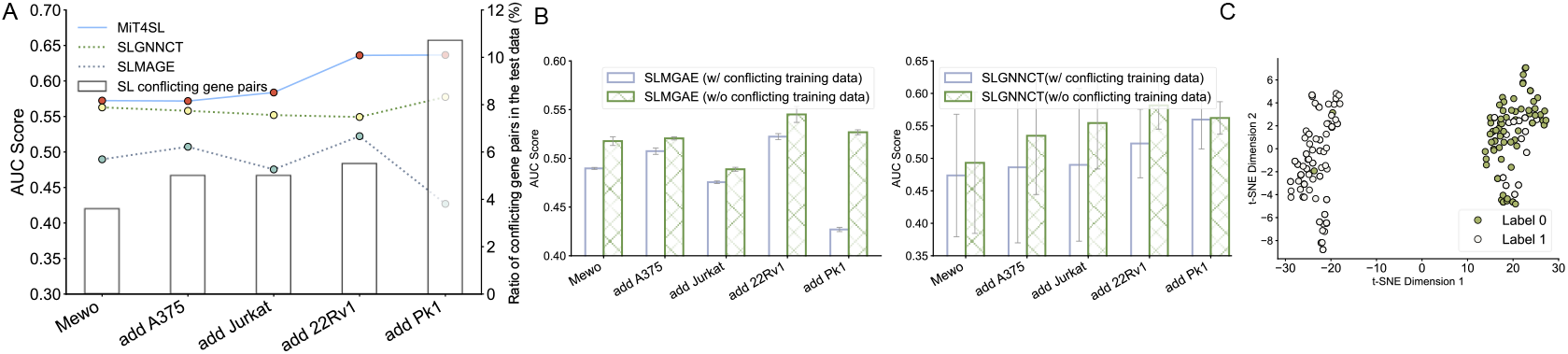
(A) Cumulative performance curve as more cell lines are added to the training set. (B) Results of SLMGAE and SLGNNCT with and without conflicting SL gene pairs in the training data for cell line-adapted SL prediction. (C) Visualization of MiT4SL embeddings for conflicting SL pairs. Gene pairs from A549 are labeled as negative (Label 0), while those from other cell lines are labeled as positive (Label 1).

We compare the performance of SLMGAE and SLGNNCT trained with and without conflicting SL labels, using the same test data, to validate this hypothesis. Both models trained without conflicting labels consistently outperform their counterparts trained with conflicting labels (Figure 5B), suggesting that such conflicts negatively impact model performance. Notably, both SLMGAE and SLGNNCT adopt gene-pair representation learning for SL prediction. As cell-line-free SL prediction methods, including but not limited to SLMGAE, they do not incorporate cell line information and instead generate universal gene-pair representations to predict SLs across all cell lines. Consequently, when SL labels for the same gene pairs vary between source and target cell lines, these methods struggle to distinguish context-specific SL interactions, leading to inaccurate predictions. SLGNNCT tends to segment the BKG based on SL gene pairs from distinct cancer types and independently learns SL relationships within each cell line. However, this segmentation strategy does not provide SLGNNCT, which relies solely on gene-pair representations, with sufficient signals to differentiate between cell lines. As a result, it remains unable to resolve conflicting SL labels and struggles to effectively leverage additional training data.

By contrast, MiT4SL benefits from incorporating additional cell lines, even when conflicting SL labels are present. This advantage arises from its triplet representation, which explicitly encodes cell line information alongside gene pair embeddings, allowing the model to dynamically adjust predictions based on cell line context. More importantly, by seamlessly integrating gene pair and cell line information, MiT4SL captures the underlying biological mechanisms driving SL relationships across different cell lines, enabling more robust SL prediction patterns. We further visualize the high-dimensional embeddings of 214 gene pairs in a 2D space using t-SNE, where MiT4SL effectively differentiates between the two classes (Figure 5C). Experimental details are provided in the Supplementary Material.

Overall, these results underscore MiT4SL’s effectiveness in addressing SL label conflicts across cell lines. By fully leveraging valuable SL label data from known cell lines, MiT4SL accurately predicts potential SL pairs in target cell lines, which holds significant importance for the utility of SL label data.

### 4.3 Case study

In addition to demonstrating the superior performance of MiT4SL through statistical metrics, we also showcase its practical utility by identifying potential SL pairs for specific cell lines. To this end, we selected two lists of predicted SL pairs by focusing on two primary genes *KRAS* and *STK11* in the A549 cell line, which is of cancer type non-small cell lung cancer (NSCLC). *KRAS*, a widely studied oncogene, has been demonstrated to have the most frequent oncogene driver mutations in the NSCLC [37]. STK11, a tumor suppressor gene, is frequently mutated in the A549 cell line [38]. The MiT4SL model was trained on SL label data from the five other cell lines: A375, Jurkat, Mewo, 22Rv1, and Pk1 (Sec. 3). Then, MiT4SL is used to predict candidate SL gene pairs. Notably, among the top 10 predictions for *KRAS*, MiT4SL successfully captures 4 well-known SL partners, including *FLT3* [39] and *JAK2* [40] (Figure 6A).

**Fig. 6.**
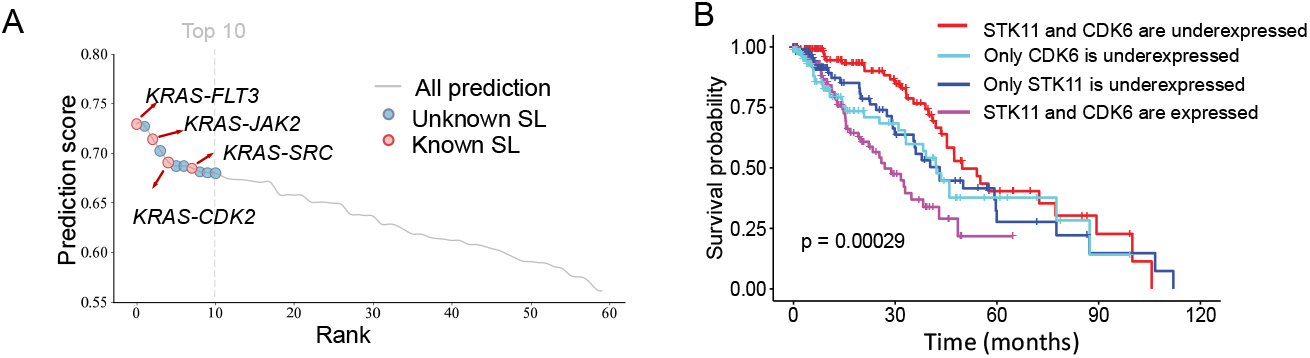
(A) Prediction scores versus rankings of candidate SL partner genes of the primary gene *KRAS* in the A549 cell line. (B) Kaplan-Meier curves and Log-rank test p-values show the survival differences based on *STK11* and *CDK6* expression levels in NSCLC patients.

For *STK11*, we assess the clinical relevance of the predicted SL interactions. Among these predictions, we find a highly ranked interaction, *STK11-CDK6*, which could represent a novel SL interaction in NSCLC. Specifically, we analyze gene expression and survival data from a total of 589 NSCLC patients derived from the TCGA database [41]. We hypothesize that co-underexpression of *STK11* and *CDK6* would increase tumor vulnerability and result in better prognosis and longer survival of the patients. Using median gene expression of each gene as a threshold, we classify the patients into four groups based on their tumor gene expression: patients with co-underexpression of both genes and patients with expression of at least one of these genes. Then, we compute the Kaplan-Meier (KM) scores and plot the corresponding KM curves for each group. A higher KM score indicates a better prognosis and longer survival of the patients. The KM score for patients with co-underexpression of both *STK11* and *CDK6* (red line) is significantly higher than other groups, with a log-rank p-value of 2.9*×* 10^−4^ (Figure 6B), indicating that the patients in this group have longer survival times. These results collectively suggest that *STK11-CDK6* is a promising and novel SL pair in NSCLC, but further wet-lab experimental validation is required to verify the relationship.

## 5 Conclusion

Synthetic lethality (SL) provides a promising strategy of cancer treatment, and it has been seen that many SL interactions are context-specific. Despite fast advancement in machine learning for SL prediction, the challenges of SL data sparsity still remain. Therefore, there is a pressing need to identify gene pairs that are most likely to be SLs in cell lines with scarce labels. Here, we propose MiT4SL, a multi-omics triplet representation learning model, which takes cell line representation as a critical component of feature embedding along with two gene embeddings. By incorporating the cell line information through explicit representation, MiT4SL effectively captures the shared biological mechanisms while addressing the conflict of SL labels across different cell lines. Consequently, MiT4SL achieves the state-of-the-art performance in the cell line-adapted scenarios. Furthermore, our case study demonstrated that MiT4SL can reliably identify novel and promising SL candidates. Thus, MiT4SL is the first solution to the critical but underexplored problem of cell line-adapted SL prediction.

## References

1. Barbara De Kegel, Niall Quinn, Nicola A Thompson, David J Adams, and Colm J Ryan. Comprehensive prediction of robust synthetic lethality between paralog pairs in cancer cell lines. Cell Systems, 12(12):1144–1159, 2021.

2. Christopher J Lord and Alan Ashworth. Parp inhibitors: the first synthetic lethal targeted therapy. Science (New York, NY), 355(6330):1152, 2017.

3. Gregory J Hannon. RNA Interference. nature, 418(6894):244–251, 2002.

4. Linsong Tang, Ronggao Chen, and Xiao Xu. Synthetic lethality: A promising therapeutic strategy for hepatocellular carcinoma. Cancer Letters, 476:120–128, 2020.

5. Yimiao Feng, Yahui Long, He Wang, Yang Ouyang, Quan Li, Min Wu, and Jie Zheng. Benchmarking machine learning methods for synthetic lethality prediction in cancer. Nature Communications, 15(1):9058, 2024.

6. Ruichu Cai, Xuexin Chen, Yuan Fang, Min Wu, and Yuexing Hao. Dual-dropout graph convolutional network for predicting synthetic lethality in human cancers. Bioinformatics, 36(16):4458–4465, 2020.

7. Zhifeng Hao, D. Wu, Yuan Fang, Min Wu, Ruichu Cai, and Xiaoli Li. Prediction of synthetic lethal interactions in human cancers using multi-view graph auto-encoder. IEEE Journal of Biomedical and Health Informatics, 25(10):4041–4051, 2021.

8. Shike Wang, Fan Xu, Yunyang Li, Jie Wang, Ke Zhang, Yong Liu, Min Wu, and Jie Zheng. Kg4sl: knowledge graph neural network for synthetic lethality prediction in human cancers. Bioinformatics, 37(S1):i418–i425, 2021.

9. Yahui Long, Min Wu, Yong Liu, Yuan Fang, Chee Keong Kwoh, Jinmiao Chen, Jiawei Luo, and Xiaoli Li. Pre-training graph neural networks for link prediction in biomedical networks. Bioinformatics, 38(8):2254–2262, 2022.

10. Shike Wang, Yimiao Feng, Xin Liu, Yong Liu, Min Wu, and Jie Zheng. NSF4SL: negative-sample-free contrastive learning for ranking synthetic lethal partner genes in human cancers. Bioinformatics, 38(S2):ii13–ii19, 2022.

11. Fangping Wan, Shuya Li, Tingzhong Tian, Yipin Lei, Dan Zhao, and Jianyang Zeng. EXP2SL: A machine learning framework for cell-line-specific synthetic lethality prediction. Frontiers in pharmacology, 11:112, 2020.

12. Kunjie Fan, Shan Tang, Birkan Gökbag, Lijun Cheng, and Lang Li. Multi-view graph convolutional network for cancer cell-specific synthetic lethality prediction. Frontiers in Genetics, 13:1103092, 2023.

13. Yasin I Tepeli, Colm Seale, and Joana P Gonçalves. ELISL: early–late integrated synthetic lethality prediction in cancer. Bioinformatics, 40(1):btad764, 2024.

14. Jingru Chen, Jianyong Pan, Yan Zhu, and Junyi Li. SLGNNCT: Synthetic lethality prediction based on knowledge graph for different cancers types. In International Conference on Intelligent Computing, pages 159–170, 2024.

15. Mahmoud Ghandi, Franklin W Huang, Judit Jané-Valbuena, Gregory V Kryukov, Christopher C Lo, E Robert McDonald III, Jordi Barretina, Ellen T Gelfand, Craig M Bielski, Haoxin Li, et al. Next-generation characterization of the cancer cell line encyclopedia. Nature, 569(7757):503–508, 2019.

16. John Paul Shen, Dongxin Zhao, Roman Sasik, Jens Luebeck, Amanda Birmingham, Ana Bojorquez-Gomez, Katherine Licon, Kristin Klepper, Daniel Pekin, Alex N Beckett, et al. Combinatorial crispr–cas9 screens for de novo mapping of genetic interactions. Nature methods, 14(6):573–576, 2017.

17. Max A Horlbeck, Albert Xu, Min Wang, Neal K Bennett, Chong Y Park, Derek Bogdanoff, Britt Adamson, Eric D Chow, Martin Kampmann, Tim R Peterson, et al. Mapping the genetic landscape of human cells. Cell, 174(4):953–967, 2018.

18. Colm J Ryan, Ilirjana Bajrami, and Christopher J Lord. Synthetic lethality and cancer–penetrance as the major barrier. Trends in cancer, 4(10):671–683, 2018.

19. Jindong Wang, Cuiling Lan, Chang Liu, Yidong Ouyang, Tao Qin, Wang Lu, Yiqiang Chen, Wenjun Zeng, and S Yu Philip. Generalizing to unseen domains: A survey on domain generalization. IEEE transactions on knowledge and data engineering, 35(8):8052–8072, 2022.

20. Mert Bulent Sariyildiz, Yannis Kalantidis, Diane Larlus, and Karteek Alahari. Concept generalization in visual representation learning. In Proceedings of the IEEE/CVF International Conference on Computer Vision, pages 9629–9639, 2021.

21. Kaiyang Zhou, Ziwei Liu, Yu Qiao, Tao Xiang, and Chen Change Loy. Domain generalization: A survey. IEEE Transactions on Pattern Analysis and Machine Intelligence, 45(4):4396–4415, 2022.

22. Xiaoqi Wang, Yingjie Cheng, Yaning Yang, Yue Yu, Fei Li, and Shaoliang Peng. Multitask joint strategies of self-supervised representation learning on biomedical networks for drug discovery. Nature Machine Intelligence, 5(4):445–456, 2023.

23. Kexin Huang, Payal Chandak, Qianwen Wang, Shreyas Havaldar, Akhil Vaid, Jure Leskovec, Girish N Nadkarni, Benjamin S Glicksberg, Nils Gehlenborg, and Marinka Zitnik. A foundation model for clinician-centered drug repurposing. Nature Medicine, pages 1–13, 2024.

24. Ziniu Hu, Yuxiao Dong, Kuansan Wang, and Yizhou Sun. Heterogeneous graph transformer. In Proceedings of the web conference 2020, pages 2704–2710, 2020.

25. Payal Chandak, Kexin Huang, and Marinka Zitnik. Building a knowledge graph to enable precision medicine. Scientific Data, 10(1):67, 2023.

26. Alexander Rives, Joshua Meier, Tom Sercu, Siddharth Goyal, Zeming Lin, Jason Liu, Demi Guo, Myle Ott, C Lawrence Zitnick, Jerry Ma, et al. Biological structure and function emerge from scaling unsupervised learning to 250 million protein sequences. Proceedings of the National Academy of Sciences, 118(15):e2016239118, 2021.

27. Amos Bairoch, Rolf Apweiler, Cathy H Wu, Winona C Barker, Brigitte Boeckmann, Serenella Ferro, Elisabeth Gasteiger, Hongzhan Huang, Rodrigo Lopez, Michele Magrane, et al. The universal protein resource (uniprot). Nucleic acids research, 33(Suppl_1):D154–D159, 2005.

28. Sam Behjati and Patrick S Tarpey. What is next generation sequencing? Archives of Disease in Childhood-Education and Practice, 98(6):236–238, 2013.

29. Kevin Titeca, Irma Lemmens, Jan Tavernier, and Sven Eyckerman. Discovering cellular protein-protein interactions: Technological strategies and opportunities. Mass spectrometry reviews, 38(1):79–111, 2019.

30. Jiacheng Lin, Hanwen Xu, Addie Woicik, Jianzhu Ma, and Sheng Wang. Pisces: A combo-wise contrastive learning approach to synergistic drug combination prediction. bioRxiv, pages 2022–11, 2022.

31. Michelle M. Li, Yepeng Huang, Marissa Sumathipala, Man Qing Liang, Alberto Valdeolivas, Ashwin N. Ananthakrishnan, Katherine Liao, Daniel Marbach, and Marinka Zitnik. Contextual AI models for single-cell protein biology. Nature Methods, 21(8):1546–1557, August 2024.

32. Guohao Li, Chenxin Xiong, Ali Thabet, and Bernard Ghanem. DeeperGCN: All you need to train deeper GCNs. arXiv preprint 2006.07739, 2020.

33. Birkan Gökbag, Shan Tang, Kunjie Fan, Lijun Cheng, Lianbo Yu, Yue Zhao, and Lang Li. SLKB: synthetic lethality knowledge base. Nucleic Acids Research, 52(D1):D1418–D1428, 2024.

34. Laurens Van der Maaten and Geoffrey Hinton. Visualizing data using t-sne. Journal of Machine Learning Research, 9(11), 2008.

35. María Luisa Menéndez, JA Pardo, L Pardo, and MC Pardo. The jensen-shannon divergence. Journal of the Franklin Institute, 334(2):307–318, 1997.

36. Zhaoxiang Cai, Sofia Apolinário, Ana R Baião, Clare Pacini, Miguel D Sousa, Susana Vinga, Roger R Reddel, Phillip J Robinson, Mathew J Garnett, Qing Zhong, et al. Synthetic augmentation of cancer cell line multi-omic datasets using unsupervised deep learning. Nature Communications, 15(1):10390, 2024.

37. Niki Karachaliou, Clara Mayo, Carlota Costa, Ignacio Magrí, Ana Gimenez-Capitan, Miguel Angel Molina-Vila, and Rafael Rosell. Kras mutations in lung cancer. Clinical lung cancer, 14(3):205–214, 2013.

38. Montserrat Sanchez-Cespedes, Paola Parrella, Manel Esteller, Shuji Nomoto, Barry Trink, James M Engles, William H Westra, James G Herman, and David Sidransky. Inactivation of lkb1/stk11 is a common event in adenocarcinomas of the lung. Cancer research, 62(13):3659–3662, 2002.

39. Hwani Ryu, Hyun-Kyung Choi, Hyo Jeong Kim, Ah-Young Kim, Jie-Young Song, Sang-Gu Hwang, Jae-Sung Kim, Da-Un Kim, Eun-Ho Kim, Joon Kim, et al. Antitumor activity of a novel tyrosine kinase inhibitor aiu2001 due to abrogation of the dna damage repair in non-small cell lung cancer cells. International journal of molecular sciences, 20(19):4728, 2019.

40. Nastaran Karimi and Seyed Javad Moghaddam. Kras-mutant lung cancer: targeting molecular and immunologic pathways, therapeutic advantages and restrictions. Cells, 12(5):749, 2023.

41. John N Weinstein, Eric A Collisson, Gordon B Mills, Kenna R Shaw, Brad A Ozenberger, Kyle Ellrott, Ilya Shmulevich, Chris Sander, and Joshua M Stuart. The cancer genome atlas pan-cancer analysis project. Nature genetics, 45(10):1113–1120, 2013.

