## Supplemental Material for "MiT4SL: multi-omics triplet representation learning for cancer cell line-adapted prediction of synthetic lethality"

### 1 Experimental settings

#### 1.1 Compared methods

We compare our MiT4SL with six state-of-the-art (SOTA) SL prediction approaches. They are briefly summarized as follows.

- SLMGAE [3] is a multi-view graph auto-encoder that predicts SL pairs by integrating the known SL graph, GO, and PPI data.
- KG4SL [9] is the first SL prediction model that encodes the genes by integrating a knowledge graph.
- PT-GNN [5] is a pre-trained GNN model to learn high-quality representations for downstream tasks, including SL prediction.
- NSF4SL [8] is a contrastive learning-based approach for SL prediction without involving any negative sample.
- MVGCN-iSL [2] is a multi-view GCN model that can predict SL pairs in a specific cell line by incorporating five cell-line-specific graphs, e.g., physical PPI network and co-expression network.
- SLGNNCT [1] leverages a graph neural network to extract features from knowledge graph to predict SL pairs for different cancer types.

Note that SLMGAE, KG4SL, PT-GNN, and NSF4SL are cell-line-free SL prediction models that ignore the genetic information of the cell lines, while MVGCN-iSL and SLGNNCT are cell-line-specific SL prediction models. To ensure a fair comparison, we implement them using the open-source code provided by the authors.

### 2 Results

#### 2.1 Performance comparison in the cell line-specific scenario

We compare MiT4SL with the SOTA baselines for SL prediction in the cell-line-specific scenario. Figure 1 shows the performance of the methods in six different cell lines (A375, A549, Jurkat, Mewo, 22Rv1, and Pk1). We adopt a 70%/10%/20% split for training, validation, and testing. Model performance is validated by 5-fold cross-validation.

As shown in Figure 1, MiT4SL achieves the best performance in the cell-line-specific scenario compared with the SOTA methods, although it is designed for the cell-line-adapted SL prediction task. First, compared to the KG4SL, PTGNN, and NSF4SL that only learn from a single modality of data, such as Biomedical knowledge graph (BKG), MiT4SL effectively utilizes multi-omics data and thus achieves significant improvements. In addition, MiT4SL also produces minor improvements over the SOTA SL prediction methods using multi-modal biological data (e.g., SLMGAE and MVGCN-iSL). There could be two reasons for this. First, multi-omics data used in MiT4SL contain rich biological information, which can enhance the generation of comprehensive and good representations. Second, two novel regularizations (i.e., $\mathcal{L}_{\text{CL}}$ and $\mathcal{L}_{\text{DLM}}$)

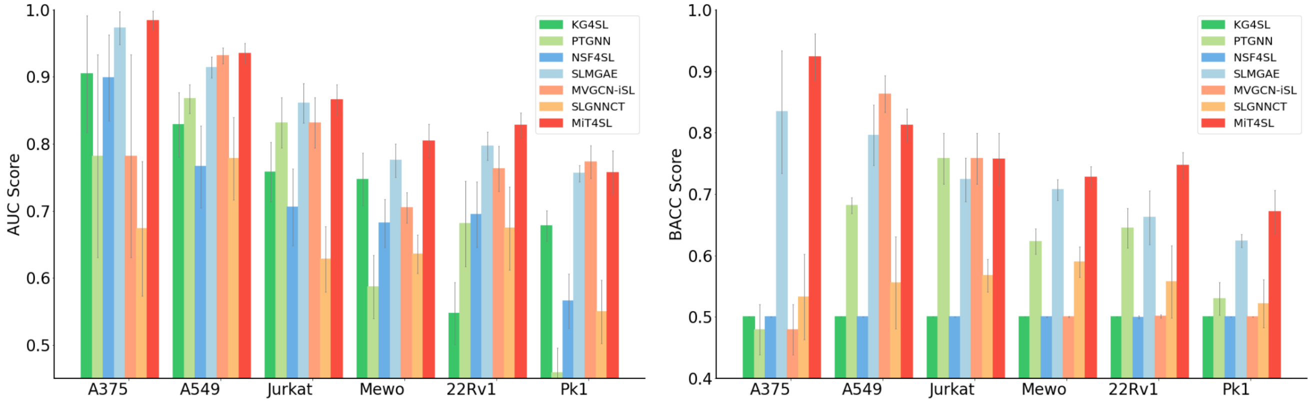


Figure 1: Performance comparison of MiT4SL and six existing approaches across six cell-lines for the cell-line-specific SL prediction task, using bar plots to represent AUC and BACC values. Each result is based on 5-fold cross validation.

proposed by our MiT4SL can effectively integrate the features from different types of data and thus improve model performance. Overall, these results confirm the superiority and effectiveness of our MiT4SL over the SOTA SL prediction methods in the cell-line-specific scenarios.

#### 2.2 Performance of MiT4SL in recommending SL partners

Results for six primary genes are provided in the Figure 2. MiT4SL achieves the highest Precision@8 across all three primary genes, demonstrating its superiority in prioritizing relevant SL candidate pairs.

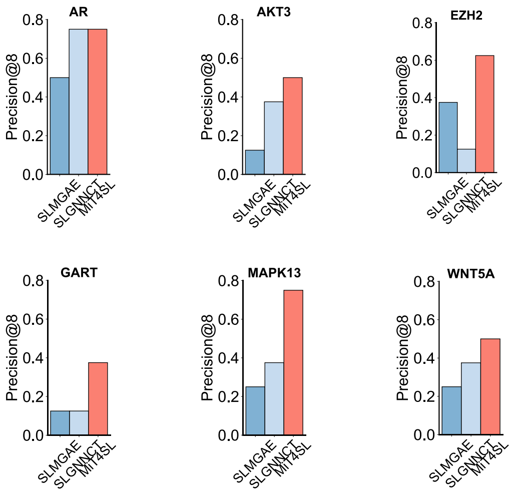


Figure 2: Comparison of MiT4SL and baseline models in recommending SL partners for primary genes in the 22Rv1 cell line. Quantitative results are evaluated using the Precision@8 score. The precision@8 score ranges from 0 to 1, where values closer to 1 indicate better performance.

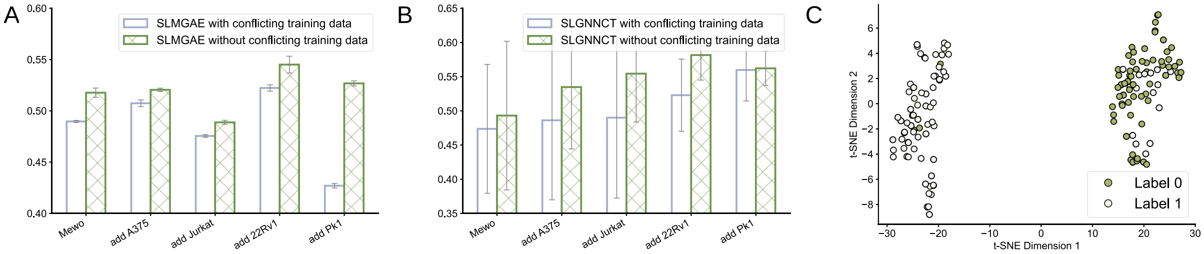


Figure 3: (A-B) Quantitative results of SLMGAE and SLGNNCT with and without conflicting SL gene pairs in the training data for cell-line-adapted SL prediction (target cell line A549). (C) Visualization of the embeddings from MiT4SL for the conflicting SL pairs. Gene pairs from the A549 cell line are labeled as negative (Label: 0), while those from other cell lines are labeled as positive (Label: 1).

Table S1: Quantitative results of ablating the multi-omics data of encoding cell lines in our MiT4SL for the cell-line-adapted SL prediction (A549 as target cell-line). '✓' means the use of the data, while '✗' refers to removing the data.

| **Cell Line Repre.** | | **Cell-line-adapted (A549 as target cell-line)** | |
|---|---|---|---|
| PPI | Prot. Seq. | AUC (%) | BACC (%) |
| ✓ | ✗ | 62.47$_{\pm 3.24}$ | 57.00$_{\pm 3.95}$ |
| ✗ | ✓ | 53.96$_{\pm 4.93}$ | 53.93$_{\pm 3.58}$ |
| ✓ | ✓ | **63.66**$_{\pm \mathbf{3.26}}$ | **58.27**$_{\pm \mathbf{2.56}}$ |

##### 2.3 Performance of MiT4SL on gene pairs with conflicting SL labels

We assess the performance of MiT4SL on these unique gene pairs with conflicting SL labels across cell lines. Briefly, we drop these gene pairs from training data (A375, Jurkat, Mewo, 22Rv1, and Pk1) and take these unique gene pairs (about 214) as test data. Then, we map the high-dimensional embeddings of these gene pairs into a 2D space by using t-SNE. MiT4SL effectively separates the embeddings of these two classes, demonstrating that triplet representation is promising in addressing these unique conflicting SL labels (Figure 3C).

#### 3 Ablation study

##### 3.1 Effect of multi-omics data in cell line representation

Our cell line representations are also extracted from the multi-omics data (PPI network and protein sequence). We performed an ablation study for the two types of biological data on the cell-line-adapted SL prediction task (A549 as target cell-line). Table S1 shows the results. Compared to single omics data (PPI only or protein sequence only), the use of multi-omics data can achieve better performance, confirming the effectiveness of multi-omics data for encoding cell lines. In addition, we observe that the PPI network containing biological semantic interactions is more important than protein sequences. This may be attributed to the rich interactions among genes, which can provide a global picture of cellular function and biological processes [4] and thus improve the model prediction performance.

##### 3.2 Effect of the cell line representation in mitigating distribution shift issue

Next, we investigate whether incorporating cell line representations can better align the predicted SL score distribution with the ideal distribution in the cell-line-specific (IID) setting, thereby mitigating distributional shift. To this end, we employ Kernel Density Estimation (KDE) [7] to visualize the predicted score distributions and use Jensen-Shannon (JS) divergence [6], denoted as $D_{\mathrm{JS}}$, as a measure of distributional similarity, where a lower indicates greater alignment with the IID setting.Taking the A549 cell line as an example, the introduction of cell line representation consistently shifts the predicted distribution closer to the IID scenario, suggesting improved generalization and reduced distribution shift (Figure 4). Specifically, in the overall prediction, incorporating cell line representation decreases $D_{\mathrm{JS}}$ from 0.7978 to

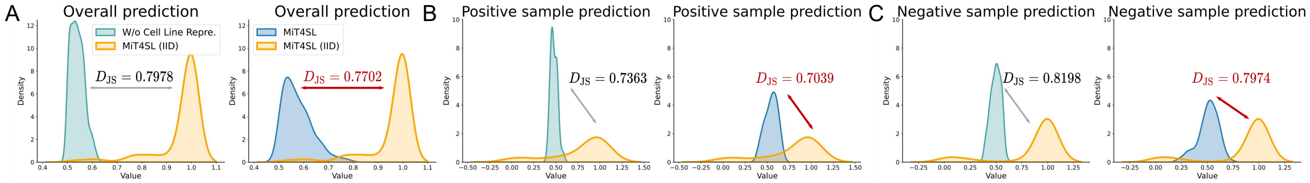


Figure 4: (A–C) Visualization of prediction distributions with and without cell line representation in cell-line-adapted scenarios, along with the prediction distribution in the MiT4SL cell-line-specific scenario (denoted as IID). $D_{\text{JS}}$ quantifies the distributional differences.

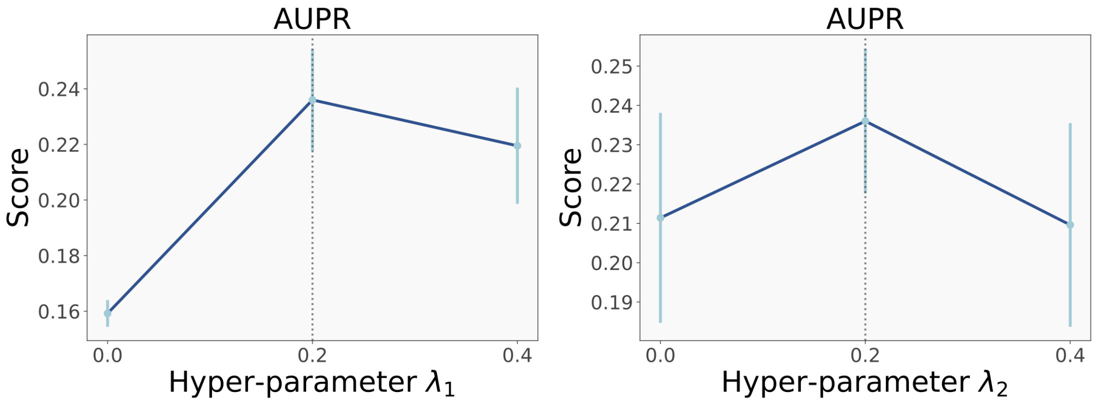


Figure 5: Quantitative results of ablating study.The results of ablating the regularizations $\mathcal{L}_{\text{CE}}$ and $\mathcal{L}_{\text{DLM}}$ in our MiT4SL model for the cell-line-adapted SL prediction task (A549 as target cell-line). Note that here $\lambda_2$ is 0.2 when ablating $\lambda_1$, while $\lambda_1$ is 0.2 when ablating $\lambda_2$.

0.7702, with similar reductions observed in the positive sample prediction ($D_{\text{JS}} = 0.7363 \rightarrow 0.7039$) and the negative sample prediction ($D_{\text{JS}} = 0.8198 \rightarrow 0.7974$). Overall, these findings highlight the importance of modeling cell line representations, as it effectively bridges the gap between real-world SL predictions and the IID assumption, ultimately reducing the distributional shifts issue in the cell-line-adapted scenarios.

##### 3.3 Effect of regularizations $\mathcal{L}_{\text{CL}}$ and $\mathcal{L}_{\text{DLM}}$

Then, we investigate the effects of the losses $\mathcal{L}_{\text{CL}}$ and $\mathcal{L}_{\text{DLM}}$ on the model performance in the cell-line-adapted scenario (A549 as target cell-line). To this end, we set different weights $\lambda_1, \lambda_2 \in \{0, 0.2, 0.4\}$ for the two losses. Additionally, when evaluating the impact of one weight, we keep the other fixed at its optimal value. We report the curves of AUC over the weights in Figure 5. First, the model performance initially improves with an increasing contribution $\lambda_1$ of the loss $\mathcal{L}_{\text{CL}}$, but later slightly degrades if its contribution keeps increasing. The best performance is obtained at $\lambda_1$= 0.2. This is because the absence of $\mathcal{L}_{\text{CL}}$ cannot remove the redundant information in the triplet representations. However, excessive bias towards it can result in the model ignoring the primary goal (i.e., predicting the SL relationship between gene pairs). Second, when we tune the weight of $\lambda_2$, it is clear that as $\lambda_2$ increases from 0 to 0.2, the performance of MiT4SL increases, which demonstrates the effectiveness of the loss $\mathcal{L}_{\text{DLM}}$. However, an excessive focus on improving modality consistency might mislead the model from its primary goal. In addition, the Hence, we set $\lambda_1 = 0.2, \lambda_2 = 0.2$.

### References

[1] J. Chen, J. Pan, Y. Zhu, and J. Li. SLGNNCT: Synthetic lethality prediction based on knowledge graph for different cancers types. In *International Conference on Intelligent Computing*, pages 159–170, 2024.

[2] K. Fan, S. Tang, B. Gökbağ, L. Cheng, and L. Li. Multi-view graph convolutional network for cancer cell-specific synthetic lethality prediction. *Frontiers in Genetics*, 13:1103092, 2023.

[3] Z. Hao, D. Wu, Y. Fang, M. Wu, R. Cai, and X. Li. Prediction of synthetic lethal interactions in human cancers using multi-view graph auto-encoder. *IEEE Journal of Biomedical and Health Informatics*, 25(10):4041–4051, 2021.

[4] G. Kar, A. Gursoy, and O. Keskin. Human cancer protein-protein interaction network: a structural perspective. *PLoS computational biology*, 5(12):e1000601, 2009.

[5] Y. Long, M. Wu, Y. Liu, Y. Fang, C. K. Kwoh, J. Chen, J. Luo, and X. Li. Pre-training graph neural networks for link prediction in biomedical networks. *Bioinformatics*, 38(8):2254–2262, 2022.

[6] M. L. Menéndez, J. Pardo, L. Pardo, and M. Pardo. The jensen-shannon divergence. *Journal of the Franklin Institute*, 334(2):307–318, 1997.

[7] G. R. Terrell and D. W. Scott. Variable kernel density estimation. *The Annals of Statistics*, pages 1236–1265, 1992.

[8] S. Wang, Y. Feng, X. Liu, Y. Liu, M. Wu, and J. Zheng. NSF4SL: negative-sample-free contrastive learning for ranking synthetic lethal partner genes in human cancers. *Bioinformatics*, 38(S2):ii13–ii19, 2022.

[9] S. Wang, F. Xu, Y. Li, J. Wang, K. Zhang, Y. Liu, M. Wu, and J. Zheng. Kg4sl: knowledge graph neural network for synthetic lethality prediction in human cancers. *Bioinformatics*, 37(S1):i418–i425, 2021.
